# Arthroscopic Raman Probe Provides Biomarkers of TMJ Tissue Composition that Account for Tissue Function

**DOI:** 10.64898/2026.10.25.761706

**Authors:** Jacob J. Giering, Erik E. Ersland, Bingyun Wang, Javier Garcia, Henry E. Brisebois, Fernando Guastaldi, Pushkar Mehra, Daniel E. Perez, David J. Chang, Mads S. Bergholt, Brian D. Snyder, Michael B. Albro

**Affiliations:** College of Engineering, Boston University, Boston, MA, USA; Department of Oral & Maxillofacial Surgery, Massachusetts General Hospital, Boston MA, USA; Department of Oral & Maxillofacial Surgery, Boston University, Boston MA, USA; Southwest Virginia Oral and Maxillofacial Surgery, Roanoke, VA, USA; Department of Oral and Maxillofacial Surgery, Tufts Medical Center, Boston, MA, USA; Centre for Craniofacial and Regenerative Biology, King’s College London, London, UK; Department of Orthopaedic Surgery, Boston Children’s Hospital, Boston, MA, USA

**Keywords:** Raman spectroscopy, temporomandibular joint, osteoarthritis, cartilage composition, biomarkers, extracellular matrix

## Abstract

**Background:** Temporomandibular joint (TMJ) disorders affect up to 25% of the population and may require diverse surgical interventions. However, intraoperative decision-making relies largely on visual assessment, which does not quantitatively characterize tissue composition or mechanical function. Raman spectroscopy offers a nondestructive approach to molecular-level tissue characterization, but its application to TMJ tissues remains unvalidated.

**Methods:** An arthroscopy-compatible Raman probe was used to characterize mandibular condyles (n=15 porcine, n=11 human) and human articular discs (n=7). Raman-derived biomarkers associated with glycosaminoglycans (GAG), collagen (COL), water (H₂O), and hydroxyapatite (HA) were evaluated against condylar indentation modulus and disc tensile modulus. Arthroscopic measurements in cadaveric TMJs assessed technical feasibility, while hyperspectral Raman imaging characterized spatial compositional distributions.

**Results:** For condylar cartilage, combined GAG and H₂O biomarkers accounted for 86% of the variation in indentation modulus (R²=0.86), compared with 4% for Outerbridge grading (R²=0.04). For articular discs, combined proline and water-associated biomarkers accounted for 88% of the variation in tensile modulus (R²=0.88). Raman spectra acquired *in situ* under simulated arthroscopic conditions were comparable in quality to *ex vivo* measurements. Raman imaging revealed distinct compositional patterns in healthy and degenerated tissues, including GAG depletion and cartilage thinning in degenerated condylar cartilage.

**Conclusions:** Raman-derived biomarkers are strongly associated with TMJ cartilage and disc mechanical properties and outperform Outerbridge grading in explaining variation in condylar indentation modulus. Arthroscopic Raman spectroscopy is technically feasible and may complement visual assessment by providing objective compositional information to support future development of tissue-informed surgical decision-making.

## INTRODUCTION

The temporomandibular joint (TMJ) is a diarthrodial joint that permits both hinge and sliding movements of the mandible, enabling mastication and speech. TMJ function depends on two specialized tissues that work in concert. First, cartilage lines the mandibular condyle and glenoid fossa, comprising a heterogeneous fibrocartilage tissue with a collagen and elastin fibril network that provides tensile and shear strength, together with a hyaline-like region enriched in negatively charged, sulfated glycosaminoglycans (GAGs)^1–3^. Second, a fibrocartilaginous articular disc contains aligned collagen fibers optimized for load distribution^4,5^. These components act synergistically to achieve the rheological and tribological properties essential for TMJ function^2,6^. Interstitial fluid pressurization supports compressive loading^6,7^, while the structural organization of the fibrocartilage and disc maintains joint stability and reduces friction during articulation^6,8,9^. The unique anatomy and regional heterogeneity of TMJ tissues—including the biconcave disc with variable tissue properties across the anterior, intermediate, and posterior zones—create a complex biomechanical system optimized for load distribution and joint stability during mastication^4^.

TMJ disorders (TMD) are painful conditions that affect up to 25% of the U.S. population and are associated with degeneration of cartilage and disc tissues^10,11^. Multiple etiologies, including trauma, internal disc derangement, joint instability, and mechanical overloading, trigger biochemical and structural breakdown of the extracellular matrix (ECM), compromising tissue material properties and joint function^10,12,13^. Disc degeneration is an important contributor to TMD progression: loss of collagen network integrity reduces the disc’s capacity to distribute loads and stabilize the joint, potentially accelerating cartilage degradation^9,14,15^. Clinical management includes physical therapy and intra-articular injections (e.g., corticosteroids, hyaluronic acid, or platelet-rich plasma) that offer temporary relief, but surgical management is often required. Surgical treatment varies from minimally invasive procedures such as arthrocentesis or arthroscopy^10,16,17^ to disc repositioning, partial or complete discectomy for internal derangement, and TMJ replacement for late-stage disease^10,18^. While these interventions have been demonstrated to be successful in specific patients, no evidence-based algorithms exist to guide surgeons in choosing which procedure may be best for treating specific TMJ pathoanatomy based on changes in tissue quality^17–19^.

MRI offers imaging of TMJ soft tissues, identifying disc displacement and deformation, and inflammatory changes in joint tissues^18,20^. However, MRI findings often do not correlate with clinical symptomatology in TMD^21,22^. Clinicians cannot reliably evaluate the quality of hard and soft tissues from clinical MRI^21,23,24^. These limitations motivated the routine use of diagnostic arthroscopy to assess TMD^16,17,24^. However, while arthroscopy allows direct visualization of joint tissues and adjacent structures, it does not provide information about the tissue composition and structure essential to tissue function. Because of these limitations, there is an unmet clinical need for a quantitative and objective diagnostic modality.

Raman spectroscopy is an inelastic light scattering technique that provides a quantitative, molecular-level ‘fingerprint’ of tissue composition by measuring vibrational modes of biochemical constituents, including molecular groups associated with proteins, sulfated GAGs, and water^25,26^. The resulting spectrum can be decomposed to estimate the relative contributions of ECM constituents without requiring labels or tissue destruction. We have recently developed an arthroscopy-compatible Raman probe that enables real-time, nondestructive assessment of cartilage composition during joint examination^27,28^. Prior validation in knee cartilage demonstrated that Raman-derived biomarkers account for 70–80% of the variation in cartilage composition and elastic modulus, significantly outperforming Outerbridge grading, quantitative MRI (T2, T2*), and histopathologic scoring^29^. Accordingly, arthroscopic Raman evaluation of TMJ tissues may offer a novel approach to objective tissue characterization (**Fig. 1A**). However, the TMJ has unique structural and biomechanical features, including specialized fibrocartilaginous discs with region-dependent properties and loading patterns distinct from other joints, necessitating tissue-specific validation of Raman biomarkers before clinical implementation.

**Figure 1:**
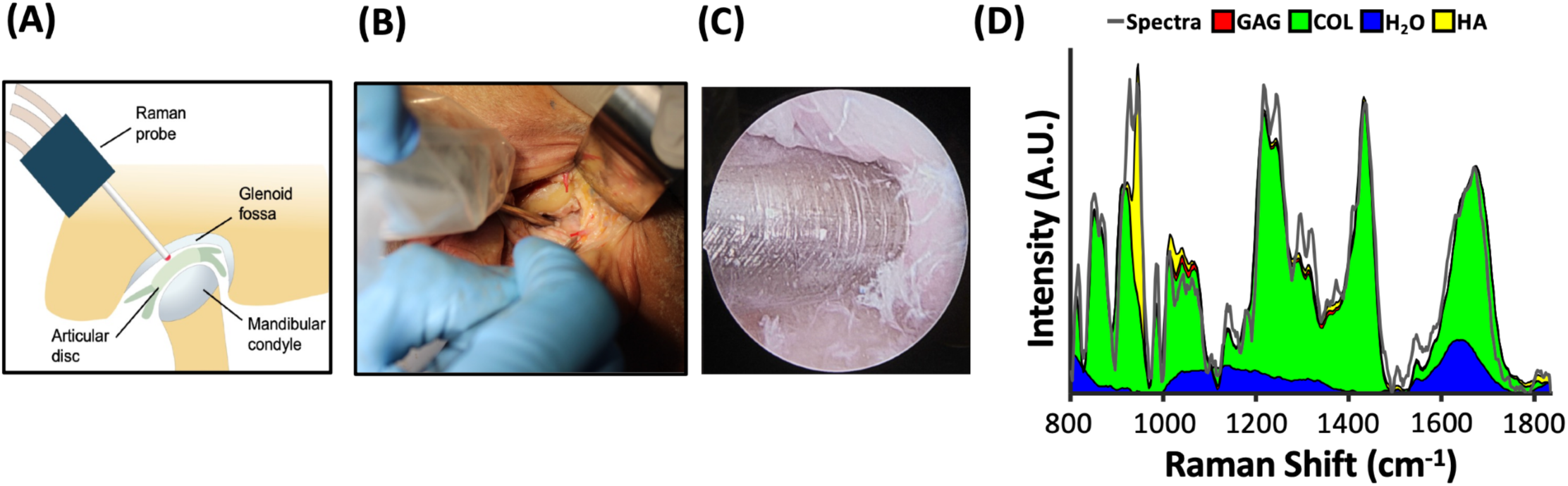
*In situ* arthroscopic Raman evaluation of the temporomandibular joint (TMJ). **(A)** Schematic of Raman probe-based arthroscopic evaluation of TMJ tissues. **(B)** Entry portal for the Raman probe during arthroscopy in a human cadaveric TMJ. **(C)** Arthroscopic Raman probe in contact with the TMJ retrodiscal tissue. **(D)** Raman spectra of retrodiscal tissue acquired arthroscopically *in situ* in a cadaver TMJ. The two-dimensional stacked area plot illustrates the cumulative contributions of glycosaminoglycans (GAG), collagen (COL), water (H_2_O), and hydroxyapatite (HA) to the measured spectra.

We hypothesize that Raman-derived biomarkers reflect compositional and structural features of TMJ cartilage and disc tissue that are associated with mechanical function across healthy and degenerated states, and that these biomarkers are more strongly associated with tissue mechanical properties than conventional arthroscopic grading. To test this hypothesis, we pursued three aims: (1) <u>TMJ tissue assessment</u>: evaluate whether Raman probe-derived compositional biomarkers account for variation in the structure and material properties of mandibular condylar cartilage and articular discs across a range of degenerative states, and whether they outperform Outerbridge grading in accounting for tissue mechanical properties; (2) <u>clinical</u> <u>feasibility</u>: demonstrate that high-quality Raman spectra can be acquired arthroscopically *in situ* from TMJ tissues in cadaveric specimens, establishing technical feasibility for clinical translation; and (3) <u>Raman imaging</u>: characterize the spatial distributions of ECM constituents in healthy and degenerated TMJ tissues to inform future refinements in probe-based spectral analysis and improve diagnostic performance.

## METHODS

### Tissue Procurement

Porcine mandibular condyle specimens (n=15) were obtained from a local abattoir. Human mandibular condyles (n=11) and articular discs (n=7) were obtained from surgical tissue discarded following TMJ arthroplasty (**Fig. 2**). All specimens were frozen at −80 °C and thawed prior to analysis. Cylindrical osteochondral cores (∅ 6 mm) were harvested from the mandibular condyles using a biopsy punch. For one human condyle and three porcine condyles, two cores were harvested per specimen, yielding a total of 18 porcine and 12 human osteochondral cores for analysis. An orthopaedic surgeon assessed condylar cartilage degeneration using the Outerbridge (OB) grading scale (grades 0–4).

**Figure 2:**
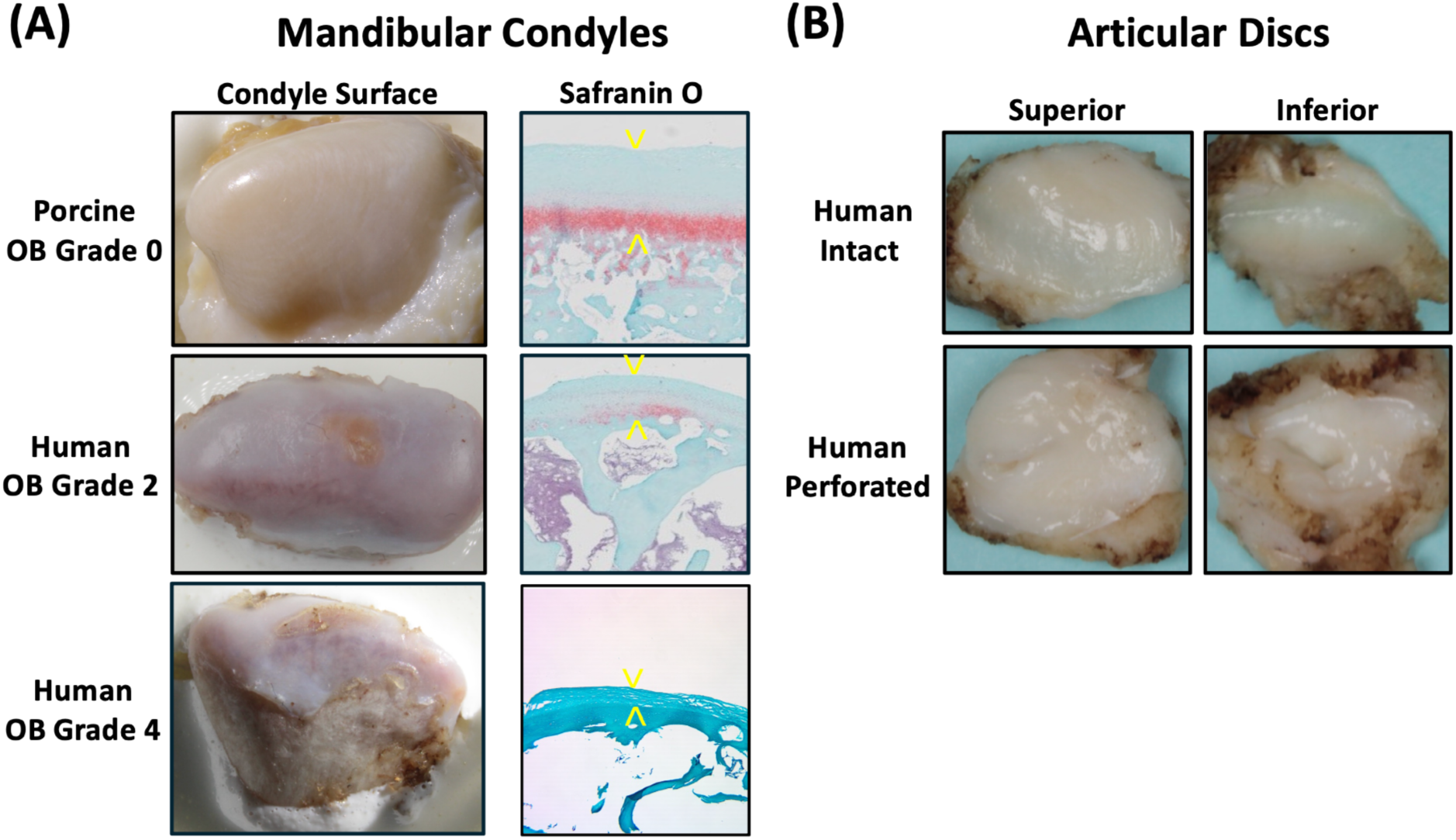
Images and histology of TMJ specimens. **(A)** Representative surface images and corresponding safranin-O/fast green-stained sections of mandibular condyle specimens with varying Outerbridge (OB) grades. Yellow arrows indicate the boundaries of the chondral layer. **(B)** Representative images of the superior and inferior surfaces of healthy (intact) and degenerated (perforated) human articular disc specimens.

Human condylar cores represented OB grades OB 0 (n=2), OB 1 (n=2), OB 2 (n=2), OB 3 (n=2), and OB 4 (n=4), while porcine cores represented OB grades OB 0 (n=15) and OB 1 (n=3). Osteochondral cores were subjected to Raman spectroscopy, indentation testing, and chondral layer thickness measurements. Chondral thickness was determined from core cross-sections using a dissecting microscope. Articular discs were sectioned into dog-bone-shaped specimens (preparation methods described below) for Raman spectroscopy and tensile testing. Five discs yielded two specimens each, one yielded one, and one yielded three (n=14 total).

### Raman Probe Instrumentation

A custom arthroscopy-compatible Raman probe (In Photonics custom model RPC785-PP-C) was used to acquire tissue spectra (**Fig. 3A**). A near-infrared diode laser (785 nm excitation wavelength, B&WTek) delivered 150 mW of excitation power through a 5.5-cm-long probe tip equipped with a sapphire ball lens (**Fig. 3A**)^27,28,30,31^. Raman-scattered light was collected by a spectrometer (Ibsen model Eagle DR-316B-LDC-DD, 0–3700 cm⁻¹, 2000 pixels). The probe was brought into gentle contact with the tissue surface, and spectra were acquired using a 30-second integration time (**Fig. 3B**). Given the exploratory nature of this study, this integration time was selected to maximize spectral signal-to-noise ratio and facilitate evaluation of the probe’s diagnostic potential.

**Figure 3:**
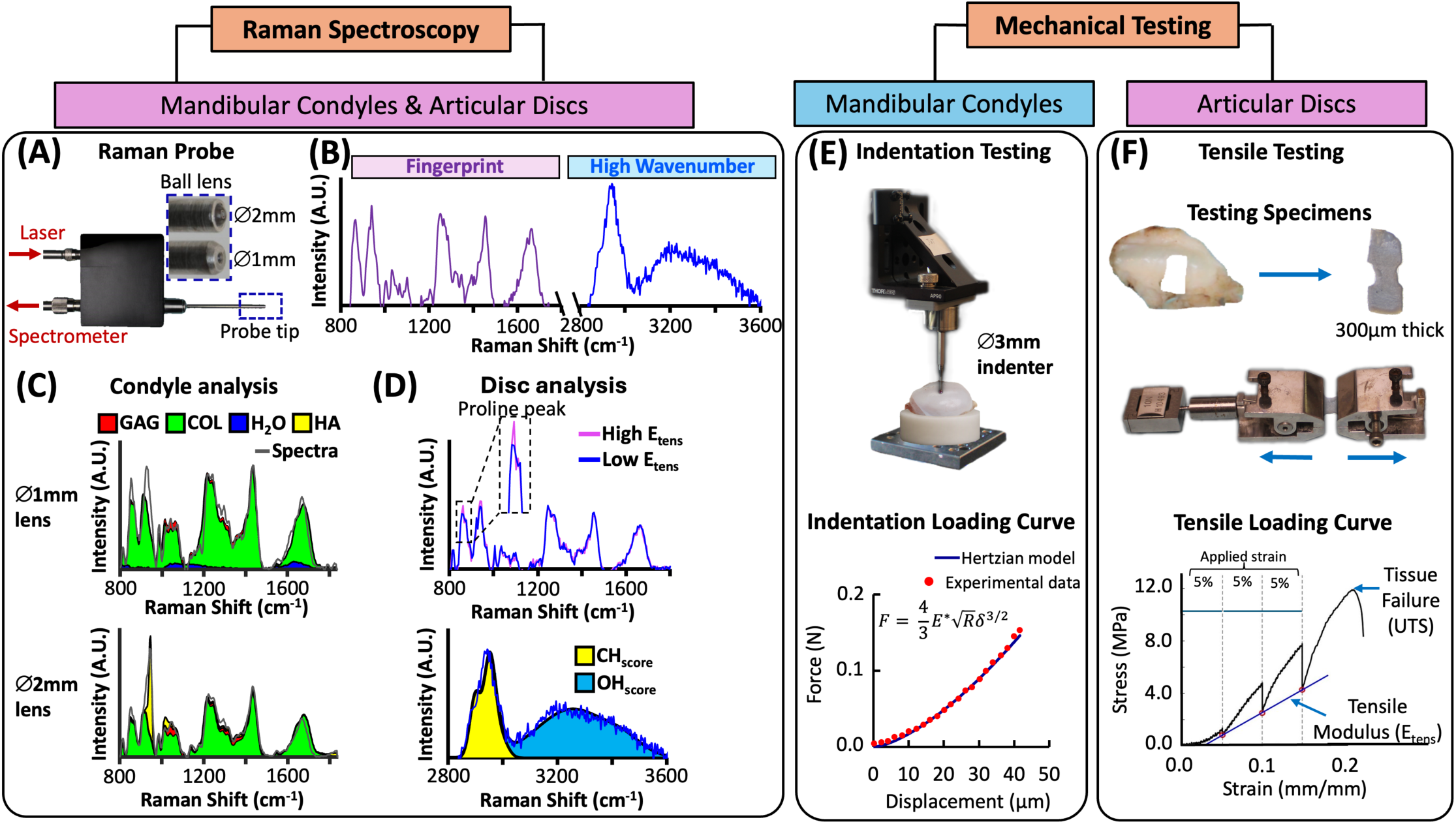
Experimental workflow for *ex vivo* Raman and mechanical analysis of condyle and disc TMJ specimens. **(A)** Arthroscopic Raman probe for intra-articular analysis of TMJ tissues. Condyle spectra are acquired via two lenses: (1) a deep-focusing ∅ 2 mm ball lens that acquires signals from subchondral hydroxyapatite, enabling estimation of chondral thickness, and (2) a shallow-focusing ∅ 1 mm ball lens that preferentially targets the chondral region, enabling evaluation of tissue composition. **(B)** Representative Raman spectra showing the fingerprint and high-wavenumber regions of tissue spectra. **(C)** <u>Condyle spectral analysis</u>: Two-dimensional stacked area plots depicting the cumulative contribution of glycosaminoglycans (GAG), collagen (COL), water (H_2_O), and hydroxyapatite (HA) to Raman spectra acquired using the ∅ 1 mm and ∅ 2 mm lenses. **(D)** <u>Disc spectral analysis</u>: The proline peak (∼856 cm⁻¹) in the fingerprint spectral region is used to calculate the Proline_score_, which reflects the stability of the collagen network’s triple-helical structure. OH peak area in high-wavenumber spectral region is used to calculate the OH_score_, which reflects tissue hydration. **(E)** <u>Condyle indentation testing</u>: Hayes-corrected Hertzian analysis of the loading curve to compute the indentation modulus (*E_ind_*). **(F)** <u>Disc tensile testing</u>: Equilibrium response to sequential 5% strain increments applied to dog-bone-shaped disc specimens, used to compute the tensile modulus (*E_tens_*).

For mandibular condyles, probes equipped with ∅ 1 mm and ∅ 2 mm ball lenses were used to interrogate different tissue depths. The ∅ 2 mm lens exhibits a depth of penetration (DOP) of approximately 470 µm, enabling detection of subchondral bone signals that provide information related to cartilage thickness. In contrast, the ∅ 1 mm lens exhibits a shallower DOP of approximately 230 µm, enabling more superficial interrogation of cartilage composition^28^. Only the ∅ 2 mm lens was used for articular disc measurements.

Raw spectra spanning 800–1800 cm⁻¹ were preprocessed using third-order polynomial baseline subtraction followed by area-under-curve normalization. The spectral profile was decomposed via reference-based multivariate linear regression using the following model:

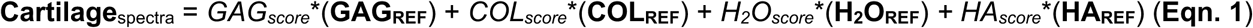

where regression coefficients (GAG_score_, COL_score_, H₂O_score_, HA_score_) represent the relative spectral contributions of the corresponding tissue constituents (**Fig. 3C**)^26,28^. Reference spectra (GAG_REF_, COL_REF_, H₂O_REF_, HA_REF_) were obtained from purified standards: chondroitin sulfate for GAG, type II collagen (chicken sternal cartilage) for collagen, phosphate-buffered saline for water, and hydroxyapatite. Each constituent score ranges from 0 to 1 and reflects its weighted contribution to the composite tissue spectrum. The Proline_score_ was calculated as the ratio of the proline-associated Raman peak near 856 cm⁻¹ to the Amide-III peak near 1245 cm⁻¹, providing a spectral marker associated with collagen structure (**Fig. 3D**)^25,32^. Spectra in the high-wavenumber region (2700– 3800 cm⁻¹) were analyzed to derive OH_score_ and CH_score_, representing spectral features associated with water and organic matrix constituents, respectively (**Fig. 3D**)^33^.

### Condyle and Disc Mechanical Testing

Mandibular condyle cores underwent indentation testing using a custom testing device equipped with a ∅ 3 mm spherical indenter. The indenter advanced 60 µm into the chondral surface at a displacement rate of 5 µm/s. The indentation modulus (*E_ind_*) was calculated by fitting the loading curve to a Hertzian contact model incorporating the Hayes correction to account for finite cartilage thickness^1,34^ (**Fig. 3E**). A Poisson’s ratio of 0.5 was assumed, consistent with the incompressible instantaneous response of articular cartilage^35,36^. Cores with cartilage thickness below 350 µm or a contact-radius-to-indentation-depth ratio greater than 1.0 were excluded from the mechanical analysis (n=5 human cores).

Rectangular explants (12 × 4 mm) were harvested from the articular discs using a leather punch, with their long axis aligned in the anteroposterior direction to follow the predominant collagen fiber orientation^14,37,38^. The explants encompassed the central intermediate zone, flanked by portions of the anterior and posterior bands. Explants were sectioned to a uniform thickness of 300 µm via Cryostat, removing the inferior and superior surfaces to obtain parallel sections and eliminate the native biconcave geometry^37,39^. The resulting sections were subpunched into dog-bone-shaped specimens (**Fig. 3F**). Tensile testing consisted of three successive 5% strain ramps, each followed by a 600-second stress relaxation period. The tensile modulus (*E_tens_*) was calculated by linear regression of the equilibrium stress-strain response^14,39^. Specimens were subsequently extended to failure at a displacement rate of 1 mm/s, and ultimate tensile strength (UTS) was calculated from the maximum engineering stress^40^.

### Cadaveric In Situ Raman Assessment

To evaluate the clinical feasibility of arthroscopic Raman assessment, Raman spectroscopy was performed *in situ* on cadaveric human TMJ tissues under simulated arthroscopic conditions (**Fig. 1B, C**). The Raman probe was introduced arthroscopically through an anterior portal, while an arthroscopic camera introduced through a posterior portal was used to visualize probe positioning. The probe tip was gently brought into contact with the articular surface of retrodiscal tissue, and spectra were acquired using a 15-second integration time. The spectral quality and the ability to resolve tissue constituents were evaluated by comparing the multivariate regression model fit (R²) of *in situ* and *ex vivo* spectra.

### Raman Imaging of TMJ Tissues

Representative healthy (porcine OB 0) and degenerated (human OB 4) condyles and discs were subjected to Raman microscopy imaging to characterize the spatial distributions of ECM constituents^41^. Tissues were bisected sagittally, and the exposed cross-sections were imaged by Raman microscopy. Hyperspectral Raman maps were acquired using a custom 785 nm Raman microscope equipped with a 60× water-immersion objective, using a spatial resolution of 10 × 10 µm and an integration time of 1.5 seconds per spectrum^42^. Semiquantitative Raman biomarkers were calculated at each pixel using the reference-based multivariate linear regression described above (**Eqn. 1**)^41^. For condyle specimens, imaging encompassed the full cartilage depth, and fingerprint spectra were decomposed to generate spatial maps of GAG_score_, COL_score_, H₂O_score_, and HA_score_. High-wavenumber spectra were additionally analyzed to generate spatial maps of CH_score_ and OH_score_.

For articular disc specimens, the regression model was modified by replacing the hydroxyapatite reference (HA_REF_) with an elastin reference (ELAST_REF_; soluble bovine elastin, Spectrum Chemical Mfg. Corp). This modification accounted for the absence of a bone-cartilage interface in the disc and the presence of elastin fibers that contribute to its mechanical resilience^5,43^. This model generated spatial maps of GAG_score_, COL_score_, H₂O_score_, and ELAST_score_ in disc specimens.

### Statistical Analysis

Regression analyses were performed to evaluate associations among Outerbridge grading, cartilage thickness, tissue mechanical properties, and Raman-derived biomarkers. Linear, polynomial, and exponential models were evaluated to characterize bivariate relationships. Statistical analyses were performed using Stata software (v18.1, StataCorp LLC, College Station, Texas).

## RESULTS

### Raman Biomarkers Correlate with Condyle Mechanical Properties and Cartilage Thickness

*E_ind_* ranged from 0.04 MPa to 0.8 MPa in human specimens and 0.1 MPa to 1.7 MPa in porcine specimens. Cartilage thickness ranged from 0.2 mm to 1.1 mm in human specimens and from 0.6 mm to 1.2 mm in porcine specimens. OB grade was not significantly associated with *E_ind_* (R^2^=0.04; p=0.33; **Fig. 4A**), although it was associated with cartilage thickness (R^2^=0.39; p<0.001; **Fig. 4B**). The Raman spectral decomposition model accounted for 85%±10% of the variation in the measured spectra. The GAG_score_ was associated with *E_ind_* (R²=0.61; p<0.001; **Fig. 5A**). Combining GAG_score_ and H_2_O_score_ strengthened the association with *E_ind_* (R²=0.86; p<0.001; **Fig. 5B**). HA_score_ was strongly associated with cartilage thickness (R²=0.80; p<0.001; **Fig. 5C**).

**Figure 4:**
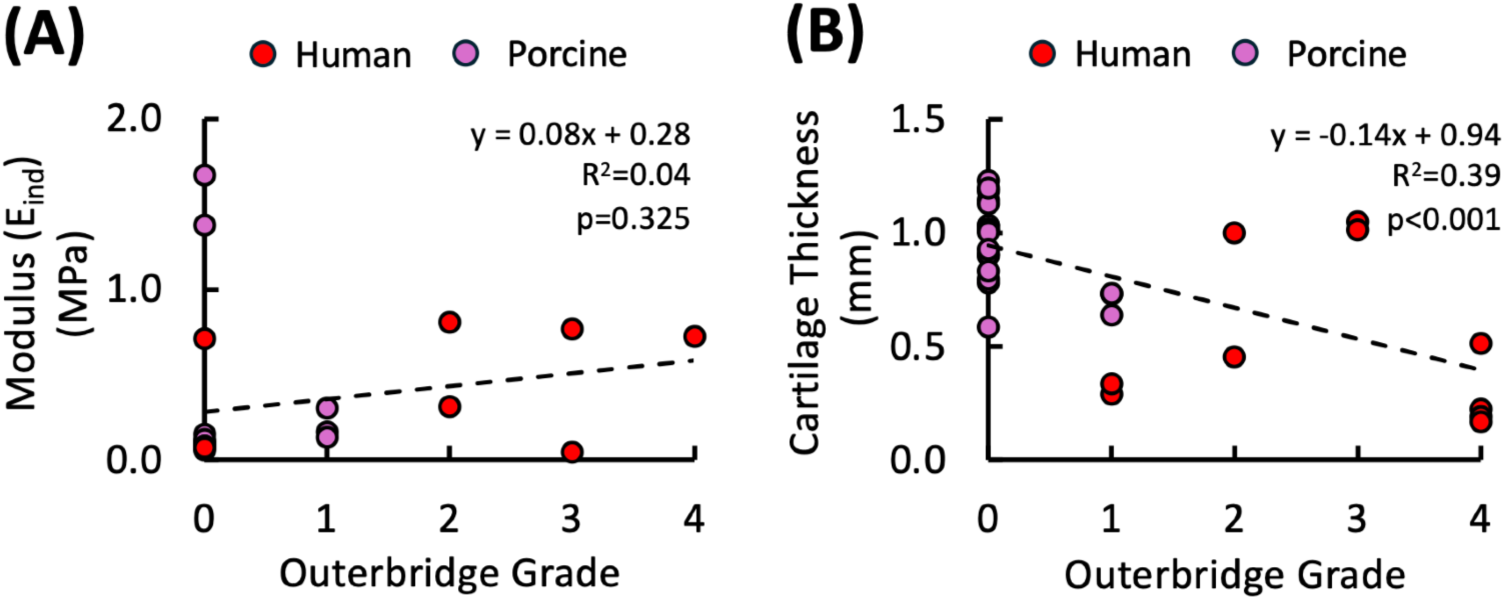
Correlation of visual grading with condyle mechanical properties and thickness. Bivariate linear regression analysis of Outerbridge grade versus **(A)** *E_ind_* and **(B)** cartilage thickness in human and porcine mandibular condyle specimens.

**Figure 5:**
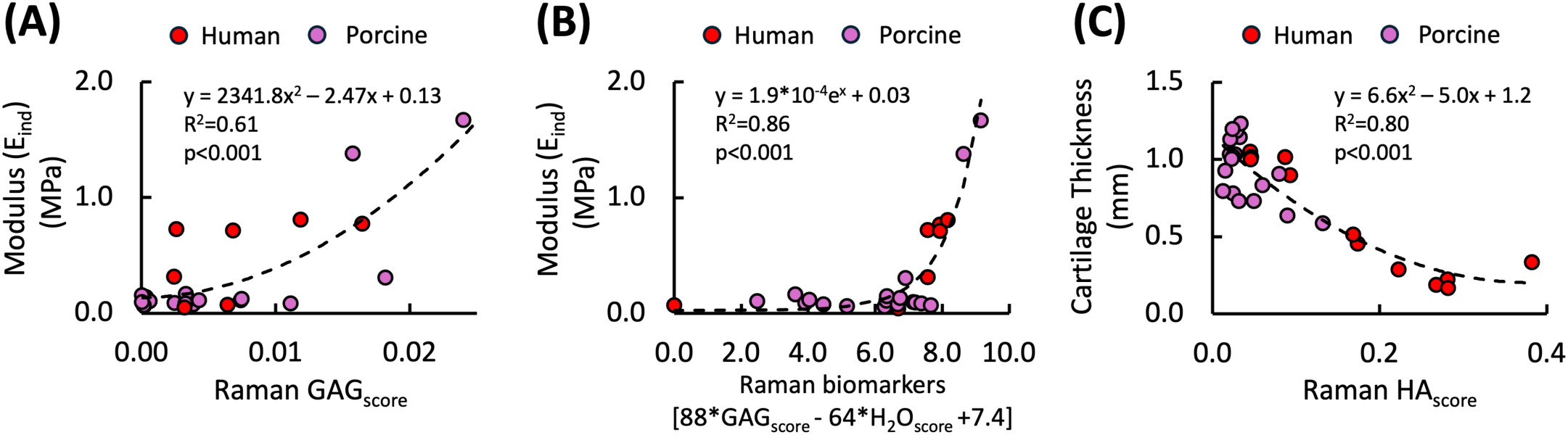
Correlation of Raman biomarkers with condyle mechanical properties and thickness. Regression analyses of *E_ind_* versus **(A)** Raman GAG_score_ and **(B)** a linear combination of GAG_score_ and H_2_O_score_ in human and porcine mandibular condyle specimens. **(C)** Regression analysis of cartilage thickness versus Raman HA_score_.

### Raman Biomarkers Correlate with Disc Tensile Properties

*E_tens_* of TMJ disc specimens ranged from ∼0.01 MPa to 10.6 MPa, and UTS ranged from 2.8 MPa to 14.7 MPa. Proline_score_ was associated with *E_tens_* (R²=0.62; p<0.01; **Fig. 6A**) and UTS (R²=0.65; p<0.01; **Fig. 6B**). OH_score_ was associated with *E_tens_* (R²=0.78; p<0.001; **Fig. 6C**), but its association with UTS did not reach statistical significance (R²=0.41; p=0.06; **Fig. 6D**). Combining Proline_score_ and OH_score_ improved associations with disc mechanical properties. The combined scores were associated with *E_tens_* (R²=0.88; p<0.001; **Fig. 6E**) and UTS (R²=0.68; p<0.001; **Fig. 6F**).

**Figure 6:**
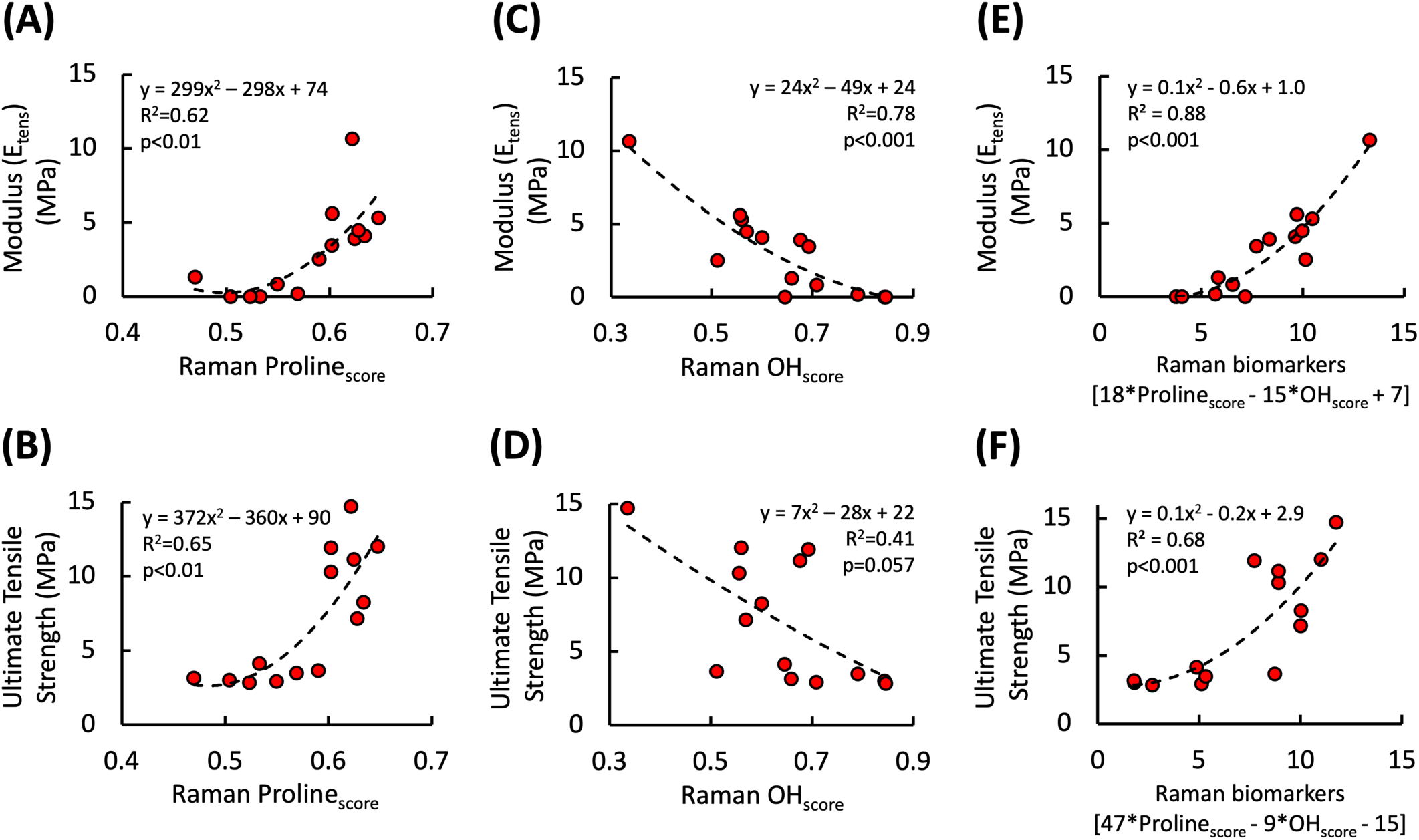
Raman biomarkers correlate with human disc mechanical properties. Regression analyses of Raman Proline_score_ versus **(A)** *E_tens_* and **(B)** ultimate tensile strength (UTS); Raman OH_score_ versus **(C)** *E_tens_* and **(D)** UTS; and a linear combination of Proline_score_ and OH_score_ versus **(E)** *E_tens_* and **(F)** UTS.

### In Situ Arthroscopic Raman Assessment

Raman spectra acquired *in situ* during simulated arthroscopy exhibited high-quality spectral decomposition (R^2^=0.86), comparable to that achieved with *ex vivo* measurements (**Fig. 1D**). These findings demonstrate the technical feasibility of arthroscopic Raman spectral acquisition from TMJ tissues under simulated clinical conditions.

### Raman Imaging Reveals Spatial Heterogeneity in TMJ Tissue Composition

Raman imaging of a healthy porcine mandibular condyle revealed spatially distinct compositional patterns across cartilage depth (**Fig. 7A**). The cartilage layer was approximately 1500 µm thick. The superficial ∼500 µm region exhibited lower GAG_score_, whereas the deeper ∼1000 µm thick region exhibited higher GAG_score_, consistent with histological characterizations of differences in matrix composition across the tissue depth^3^. HA_score_ increased near the osteochondral interface, consistent with the presence of hydroxyapatite in the underlying mineralized tissue. In contrast, the degenerated human condyle exhibited marked cartilage thinning and near-complete depletion of the GAG-associated Raman signal. These spatial patterns illustrate the ability of Raman imaging to resolve compositional differences between healthy and degenerated condylar cartilage.

**Figure 7:**
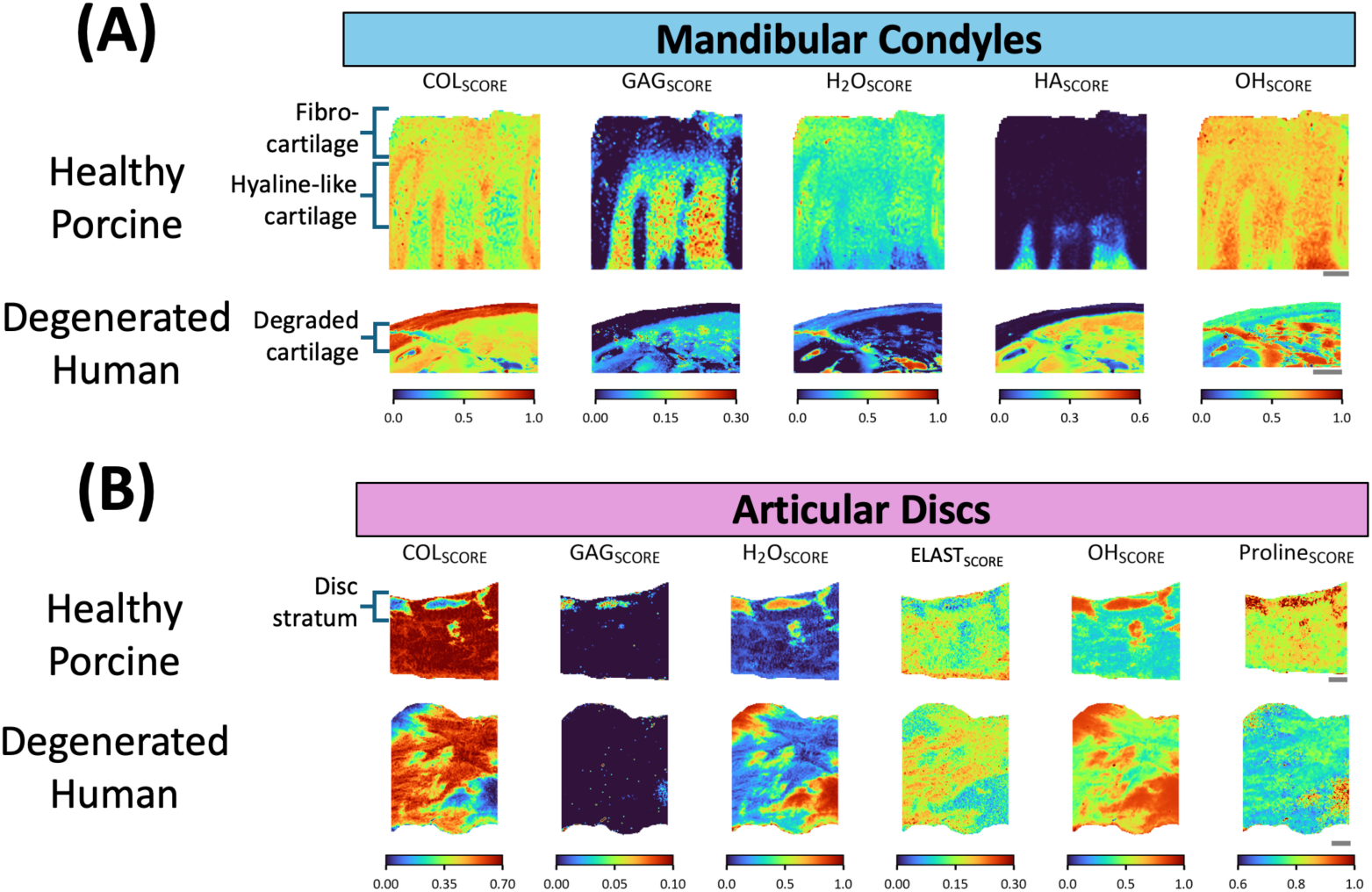
Raman microscopy-derived compositional maps of TMJ condyles and discs. **(A)** Mandibular condyle maps from healthy porcine (OB 0) and degenerated human (OB 4) specimens illustrating the spatial distribution of COL_score_, GAG_score_, H_2_O_score_, HA_score_, and OH_score_. **(B)** Articular disc maps from healthy porcine and degenerated human specimens illustrating the spatial distribution of COL_score_, GAG_score_, H_2_O_score_, ELAST_score_, OH_score_, and Proline_score_. Scale bars: 250 µm.

Raman imaging of the healthy porcine disc revealed relatively uniform collagen-associated signal, with localized regions of elevated GAG_score_ and OH_score_. In the degenerated human disc, the spatially averaged COL_score_ was lower than in the healthy porcine disc (0.45 versus 0.63), whereas the average OH_score_ was higher (0.60 versus 0.41). The average ELAST_score_ was similar between the degenerated human and healthy porcine discs (0.14 versus 0.15). The Proline_score_ map also indicated reduced proline-associated signal in the degenerated disc (**Fig. 7B**). Together, these findings demonstrate spatial differences in Raman-derived compositional biomarkers between the healthy porcine and degenerated human TMJ tissues.

## DISCUSSION

This is the first study demonstrating the technical feasibility of arthroscopic Raman spectroscopy for characterizing TMJ tissues and establishing associations between Raman-derived biomarkers and tissue mechanical properties. Multivariate decomposition of Raman spectra quantifies the relative contribution of major ECM constituents (GAG_score_, COL_score_, H₂O_score_, HA_score_) that contribute to the compositional integrity and material properties of cartilage and disc tissues essential for joint function. For condylar cartilage, Raman-derived biomarkers reflect compositional features associated with GAG-mediated Donnan osmotic swelling pressure and interstitial fluid load support, which contribute to compressive function^3,6^. The combination of GAG_score_ and H₂O_score_ was associated with 86% of the variation in *E_ind_*, demonstrating a strong relationship between Raman-derived compositional biomarkers and cartilage mechanical properties. Additionally, HA_score_ was associated with 80% of the variation in cartilage thickness, suggesting its potential to characterize structural erosion of the chondral layer. For articular discs, Raman biomarkers capture collagen-associated spectral features and tissue hydration, which are relevant to tensile and shear resistance during mastication. Specifically, Proline_score_ (a spectral marker associated with collagen structure) and OH_score_ (a water-associated spectral marker) together accounted for 88% of the variation in *E_tens_* in the fitted regression model, highlighting the potential of Raman-derived biomarkers to characterize disc mechanical properties.

Raman-derived biomarkers showed stronger associations with condylar cartilage mechanical properties than conventional Outerbridge grading. Outerbridge grading was not significantly associated with *E_ind_* (R^2^=0.04) and showed a moderate association with cartilage thickness (R^2^=0.39), demonstrating that macroscopic visual assessment provides limited information about the compositional and structural features associated with tissue function. In comparison, Raman biomarkers accounted for 86% of the variation in *E_ind_*, indicating that composition-based assessment may provide substantially more information about cartilage mechanical properties than visual grading alone. These findings support Raman spectroscopy as a promising approach for objective, intraoperative assessment of TMJ tissue composition and its relationship to mechanical function.

The acquisition of Raman spectra *in situ* from cadaveric specimens under simulated arthroscopic conditions demonstrates the technical feasibility of integrating probe-based measurements into arthroscopic workflows. Quantitative assessment of tissue composition could ultimately complement surgical evaluation and inform evidence-based decision-making. For example, if TMJ tissues are functionally competent, disc preservation or reconstruction may be appropriate. If disc tissue is functionally incompetent but cartilage remains competent, clinicians may consider targeted discectomy. If both tissues are functionally incompetent, more invasive procedures such as arthroplasty may be necessary. Clinical use would require prospective studies to establish biomarker thresholds and show that Raman-guided decisions improve patient outcomes. A composition-guided approach could complement current practice, in which treatment selection is informed by clinical presentation, imaging, surgical findings, and surgeon judgment.

Hyperspectral Raman imaging provided spatially resolved maps of ECM-associated spectral features and enabled comparison of these distributions with probe-derived biomarkers. Imaging of healthy porcine and degenerated human condyle and disc specimens demonstrated distinct spatial patterns in Raman-derived compositional scores. Healthy porcine condylar tissue exhibited a relatively thick cartilage layer with elevated GAG_score_ in deeper regions, whereas the degenerated human specimen showed cartilage thinning and near-complete depletion of the GAG-associated Raman signal. Similarly, disc imaging revealed differences in collagen-associated and water-associated signals between the healthy porcine and degenerated human specimens. Because these comparisons involved different species and only a small number of specimens, the findings are preliminary and do not independently validate the probe-derived biomarkers. Beyond providing spatial information, imaging identifies tissue-specific ECM constituents not yet incorporated into probe analysis. Notably, elastin, a structural component of the TMJ disc that contributes to tissue elasticity and resilience^5,43^, was quantified in imaging but not yet integrated into probe-based models. Incorporating an elastin-associated metric into probe analysis may improve the characterization of disc mechanical properties by reflecting viscoelastic tissue behavior. Additionally, hyperspectral imaging can investigate other matrix constituents, including proteoglycan subtypes and collagen crosslinks, that may contribute to regional tissue heterogeneity^25^. These analyses could inform future refinements of probe-based biomarkers. Although imaging of healthy and degenerated specimens was preliminary, this proof-of-concept establishes a workflow for evaluating spatial ECM heterogeneity and refining multivariate models in larger cohorts.

A limitation of this study is the exclusive use of human specimens from arthroplasty donors, precluding direct comparison with healthy human tissue. To include tissue across a spectrum of degenerative states, we supplemented the analysis with macroscopically normal porcine condylar cartilage. However, this interspecies comparison does not account for potential differences in TMJ tissue composition, structure, and material properties between species. While porcine TMJ anatomy and biomechanics are well-characterized and frequently serve as relevant preclinical models^37,39,44^, species-specific variations may influence the generalizability of structure–function relationships to human tissue. Future studies using cadaveric specimens across a range of degenerative states will enable direct evaluation of Raman biomarkers in healthy and diseased human TMJ tissues and help establish clinically relevant diagnostic thresholds.

An additional limitation is that the mechanical assays employed in this study do not fully capture the complex, multiaxial loading environment of TMJ tissues *in vivo*. While indentation testing for condylar cartilage and uniaxial tensile testing for discs are standard methods for characterizing tissue mechanical function, they do not account for the combined compressive, shear, and tensile stresses these tissues experience during mastication^11,45^. Future work could use finite element modeling that incorporates sample-specific tissue composition and structure derived from Raman biomarkers to simulate physiologically relevant loading conditions and further evaluate the relationships between tissue composition, structure, and mechanical function. These findings support further development of Raman spectroscopy as an objective, minimally invasive diagnostic adjunct to TMJ arthroscopy. Real-time, intraoperative assessment of tissue composition could ultimately inform patient-specific treatment strategies by enabling more objective evaluation of tissue quality and supporting the selection of therapeutic interventions. Beyond diagnostics, Raman-derived biomarkers may provide standardized, objective outcome measures for evaluating the efficacy of chondroprotective and chondroregenerative therapies in clinical trials. Future studies in larger human cohorts and longitudinal animal models are needed to establish diagnostic thresholds, validate the relationship between Raman biomarkers and tissue function, and determine whether Raman-guided assessment improves clinical decision-making and patient outcomes.

## ACKNOWLEDGEMENTS

Research reported in this publication was supported by the National Institute of Arthritis and Musculoskeletal and Skin Diseases under award number R01AR081393, the Arthritis Foundation, MTF Biologics, and the Boston University Ignition Award. The opinions, findings, and conclusions or recommendations expressed are those of the authors and do not necessarily reflect the views of the National Institutes of Health.

